# Sensory innervation of masseter, temporal and lateral pterygoid muscles in common marmosets

**DOI:** 10.1101/2023.02.10.528062

**Authors:** Anahit H. Hovhannisyan, Karen A. Lindquist, Sergei Belugin, Jennifer Mecklenburg, Tarek Ibrahim, Meilinn Tram, Tatiana M. Corey, Adam B. Salmon, Daniel Perez, Shivani Ruparel, Armen N. Akopian

**Author notes:** Corresponding authors: Armen N. Akopian The School of Dentistry, University of Texas Health Science Center @ San Antonio 7703 Floyd Curl Drive, San Antonio, TX 78229-3900.

## Abstract

Myogenous temporomandibular disorders (TMDM) is associated with an increased responsiveness of nerves innervating the masseter (MM), temporal (TM), medial pterygoid (MPM) and lateral pterygoid muscles (LPM). This study aimed to examine sensory nerve types innervating MM, TM and LPM of adult non-human primate - common marmosets. Sensory nerves are localized in specific regions of these muscles. Pgp9.5, marker for all nerves, and NFH, a marker for A-fibers, showed that masticatory muscles were predominantly innervated with A-fibers. The proportion of C- to A-fibers was highest in LPM, and minimal (6-8%) in MM. All C-fibers (pgp9.5^+^/NFH^-^) observed in masticatory muscles were peptidergic (CGRP^+^) and lacked mrgprD, trpV1 and CHRNA3, a silent nociceptive marker. All fibers in masticatory muscles were labeled with GFAP^+^, a myelin sheath marker. There were substantially more peptidergic A-fibers (CGRP^+^/NFH^+^) in TM and LPM compared to MM. Almost all A-fibers in MM expressed trkC, with some of them having trkB and parvalbumin. In contrast, a lesser number of TM and LPM nerves expressed trkC, and lacked trkB. Tyrosine hydroxylase antibodies, which did not label TG, highlighted sympathetic fibers around blood vessels of the masticatory muscles. Overall, masticatory muscle types of marmosets have distinct and different innervation patterns.

## Introduction

Myogenous temporomandibular disorders (TMDM) are the most prevalent group of painful orofacial conditions^1-3^. Among musculoskeletal chronic pain conditions, TMDM is the second most widespread after chronic low back pain^4^. Unlike well-localized cutaneous pain, TMDM is often manifested as referred pain to other deep tissues (i.e. toothache, headache)^5^. Current knowledge on TMDM related pain mechanisms is limited and further understanding is confounded by conflicting evidence concerning changes in superficial sensitivity seen in patients with craniofacial myalgia^6^. Thus, TMDM is often not accompanied by such clinical signs as histopathologic evidence of injury or inflammation^7^. Therefore, some studies^8^ have classified TMDM as nociplastic pain^9^. Despite these debatable points, there is an agreement that TMDM leads to sensitization of nociceptive and non-nociceptive trigeminal ganglia (TG) sensory neurons innervating the masticatory muscles, leading to increase signal input into the central nervous system^10^.

Increased responsiveness of sensory neurons during TMDM could occur in any muscle type controlling temporomandibular joint (TMJ) articulation, including superficial and in the deep heads of masseter muscle (MM), temporal muscle (TM), medial pterygoid closing muscles (MPM) as well as gliding superior and inferior heads of the lateral pterygoid muscles (LPM) ^11,12^ (*Fig 1*). These masticatory muscles are innervated by the masseteric nerve (MN) for MM and LPM, the auriculotemporal nerve (ATN) for TMJ and maybe LPM, and temporal nerve (TN) for TM (*Fig 1*). ATN plays critical role in the pathophysiology of TMJD^13^, while the MN, posterior deep TN and LPM are sensitized during TMDM^14^ (*Fig 1*).

**Figure 1.**
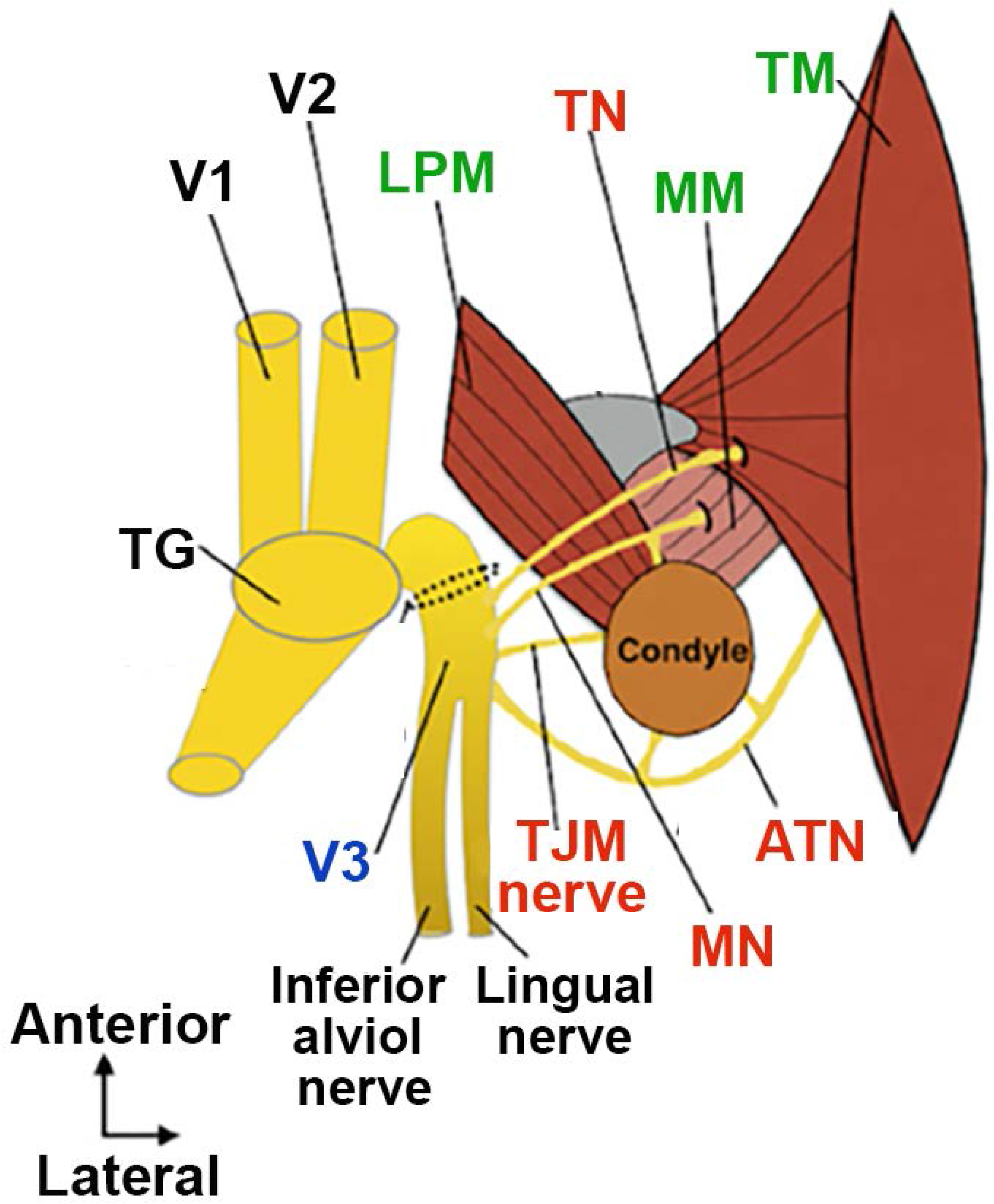
Schematic for sensory neuronal innervation of masticatory muscles. V1 – ophthalmic nerve; V2 – maxillary nerve; V3 – mandibular nerve; MN – masseteric nerve; ATN - auriculotemporal nerve; TN – posterior deep temporal nerve; TJM nerve – TJM branch of mandibular nerve; LPM - lateral pterygoid muscle; MM – masseter muscle; TM – temporal muscle.

Unlike the inferior alveolar nerve^15,16^, ATN, MN, TN and TMJ nerves are almost uncharacterized and there is only scant information on the types of afferent fibers and their function and plasticity within these branches of the mandibular (V3) nerve. The existing data was generated via extracellular recordings from the submandibular and buccal regions, which are innervated by the ATN. Recordings indicate that ATN may contain C-fibers and slow adapting A-fibers^17-19^. Electrophysiological characterization and immunohistochemistry with sensory nerve fiber markers showed that mouse MM is almost exclusively innervated by myelinated fibers^20^. To properly understand and treat TMDM pain further knowledge of the type of peripheral sensory innervation in the masticatory muscles is crucial.

We and others have demonstrated that expression patterns and characteristics of dorsal root (DRG) and TG sensory neurons depends on innervation targets ^20-22^. Moreover, there is significant differences in transcriptomic profiles of sensory neurons between rodents and humans ^23-25^. These critical differences could explain to some extent why translation of findings in rodents to clinical settings has been challenging, and stresses the need to further investigate sensory systems in other species that might better model the human anatomy and function. In this respect, nonhuman primate (NHP) models are of particular interest for additional pre-clinical testing towards translation. In this study, we used common marmosets (Callithrix jacchus), a well-characterized new world monkey used commonly for research across multiple fields including toxicology, neurological diseases, reproductive biology and aging. We report on immunohistochemistry (IHC) and known molecular markers of sensory nerve types used to characterize and identify the neuroanatomical distributions of these fibers in MM, TM and LPM of naïve adult common marmosets.

## Results

### Localization of sensory nerves in masticatory muscle

Innervation of skin of limbs in animals and humas has even distribution of sensory nerve fibers^26-28^. An innervation pattern for masticatory muscle could be uneven^29^. Thus, mouse MM innervation is localized along route of the MN trunk^20^. Locations of main MN trunk and branches within MM and LPM, and main TN trunk within TM are schematically shown in *Figure 2*. We evaluated distribution of all sensory fibers and A-fibers in male marmoset masticatory muscles. A-fibers were labeled with NFH^30,31^ and all sensory fibers with pgp9.5^31,32^. *Figure 2 upper panels* show that pgp9.5^+^ and NFH^+^ nerves are distributed within a particular area of MM of marmosets. These nerve distribution patterns were similar to those observed in mice^20^, in which nerve ends were located along a line between deep and superficial portions of MM. TM innervation was also highly localized and was detected mainly in an anterior portion of TM (*Figure 2 middle panels*). Accordingly, TM was heavily innervated in TM portion connected to the tendon, which is extended to mandible. LPM were innervated along MN trunk traveling toward condyle/TMJ (*Figure 2 bottom panels*). Overall, masticatory muscles of marmosets had localized sensory nerve innervation along main trunks of MN and TN.

**Figure 2.**
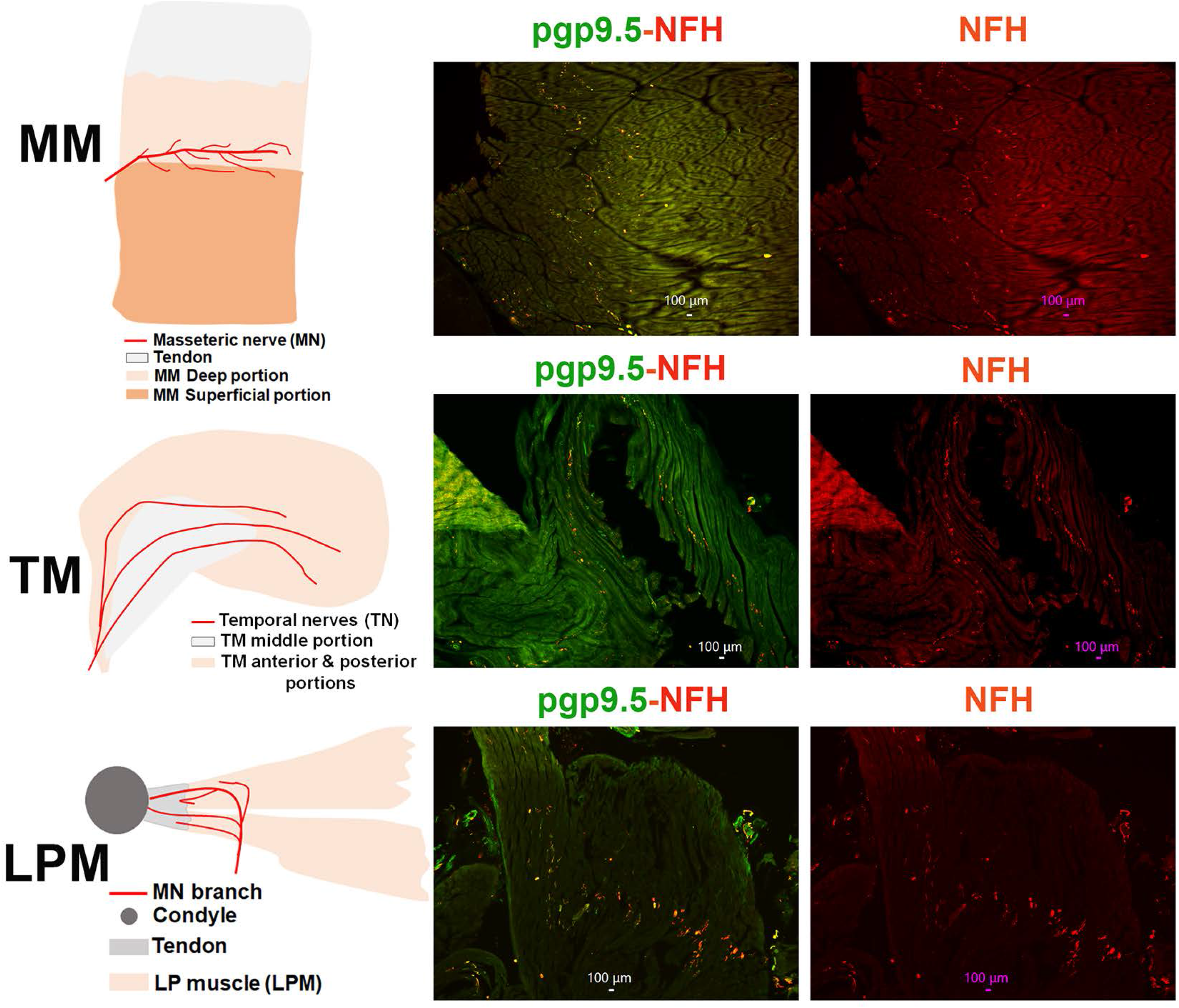
Location of pgp9.5 and NFH-positive fibers in MM, TM and LPM of adult male marmosets. *Left column* shows schematic for sensory nerves in marked/specified masticatory muscles. *Middle column* shows expression of pgp9.5 and NFH-positive fibers in MM, TM and LPM. Right column shows expression of NFH-positive fibers in MM, TM and LPM. Pictures from MM, TM and LPM as well as antibodies used and corresponding colors are indicated. Scales are presented in each microphotograph.

### Detections and quantifications of sensory nerves in masticatory muscles of marmosets

There are several approaches to detect and quantify fibers. These approaches have stronger and weaker points, and applied depending on aims and tool availability. Thus, animal reporter lines cannot be applied for marmosets as they are used in mice. Hence, sensory fibers are detected using cryosections and validated antibodies, which does not have autofluorescence and produce expression patterns in marmoset trigeminal ganglion (TG) neurons. Antibodies used in these studies were validated this way using TG labeling and auto-fluorescent validation of secondary antibodies as described before^20,33^.

Image J provides several approaches to quantify nerve fiber length^34,35^. This approach is useful for studies on neuronal morphology and neurite outgrowth. The main drawback of this approach is that the length of individual fibers can vary with each section due to unequal distribution of fibers throughout the depth of the tissue. Measuring length of fibers is not an appropriate approach for our study, since it is almost exclusively used to quantify nerve degeneration or regeneration processes, which are outside of scope of this study. Another approach assesses density of fibers using Image J platform^36,37^. This approach is required even distribution of fibers through tissues, which is not the case for masticatory muscle (*Fig 2*).

For this study, we use the manual quantification of fiber numbers, which is extensively used to assess neuropathies in animal models^38^ and clinic^39^. This approach selection were made for several reasons: a) uneven distribution of fibers in masticatory muscles; b) this study does not aim investigating regeneration or degeneration; c) it is used for neuropathy assessments, since provided for greater control over the process especially when identifying nerve fibers from artifact or debris and d) for this particular study, quantification of fiber density and length does not provide extra information on type of fibers innervating masticatory muscles.

### Expression of C- and A-fibers in the masticatory muscles

To examine the proportion of C-fibers compared to A-fibers in male marmoset masticatory muscles, we labeled A-fibers with NFH^30,31^ and all sensory fibers with pgp9.5^31,32^. Pgp9.5^+^ and NFH^+^ fibers were detected at the junction of superficial and deep heads of MM (*Fig 2*). Similar innervation pattern was observed in mouse MM^20^. Both Pgp9.5^+^ and NFH^+^ fibers had localized distribution in TM and LPM as well (*Fig 2*). Moreover, superior and inferior LPM had similar distribution patterns for Pgp9.5^+^ and NFH^+^ fibers (*Fig 2*).

We measured proportion of different fibers relatively to NFH, since unlike NFH antibodies produced in chickens, a majority of antibodies, including pgp9.5, were produced in rabbits. Alternative pgp9.5 monoclonal antibodies generated in mice and reported previously^36,37^ showed strong non-specific secondary antibody-independent labeling and high auto-fluorescence. NFH and pgp9.5 labeling overlapped in a majority of sensory nerves in MM (*Fig 3*). TM and LPM had substantially more proportion of C-fibers (pgp9.5^+^/NFH^-^) compared to MM (*Figs 3, 4A*; 1-way ANOVA; F (2, 4) = 44.48; P=0.002; n=2-3). Thus, we estimated that MM had ≈8% C-fibers and 92% A-fibers, while TM and LPM had ≈30-35% C-fibers and 65-70% A-fibers (*Fig 4A*). Overall, masticatory muscles were found to be predominantly innervated by A-fibers. MM contained the highest proportion of A-fibers, whereas LPM had lowest.

**Figure 3.**
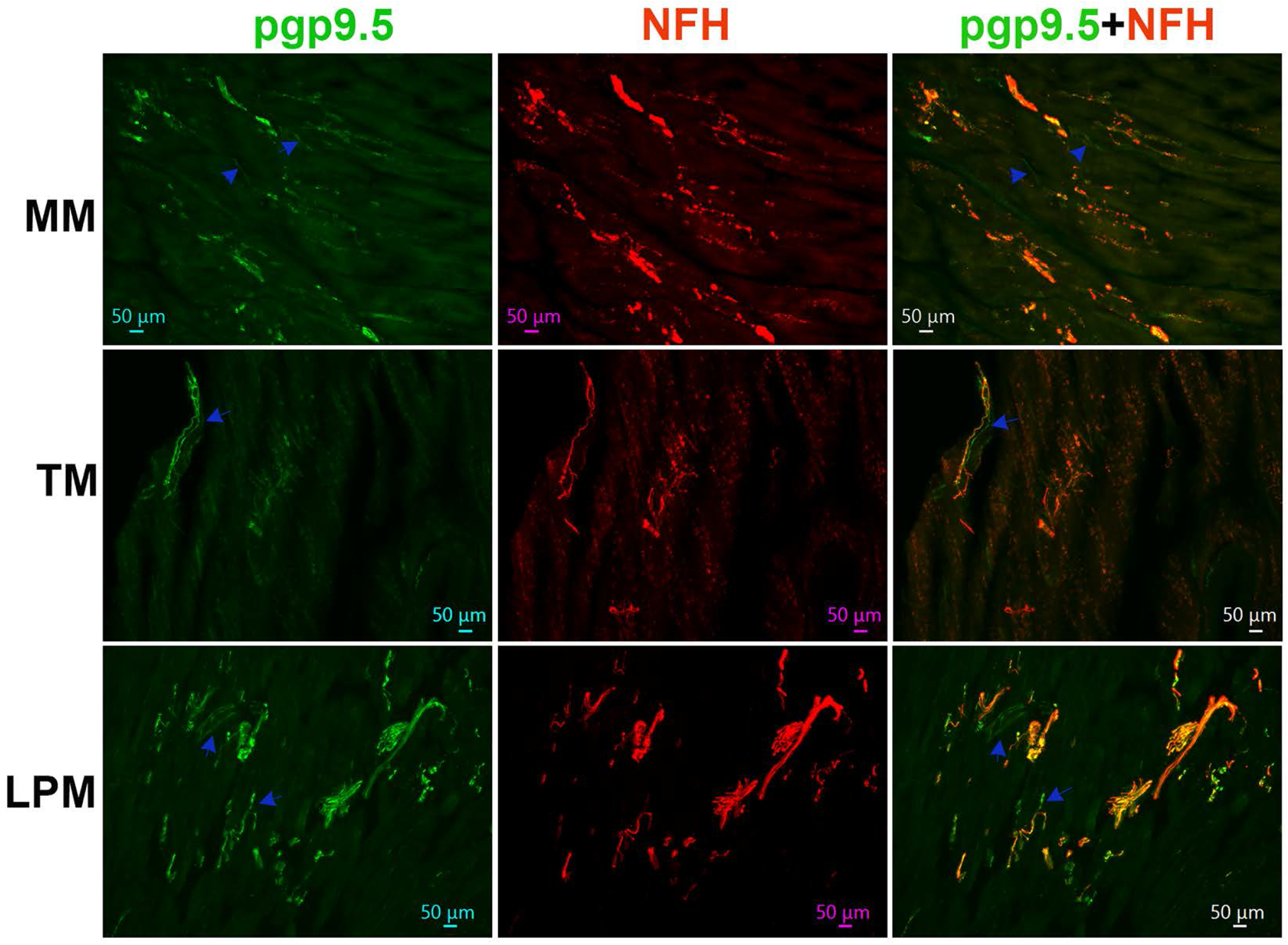
Representation of pgp9.5 and NFH-positive fibers in MM, TM and LPM of adult male marmosets. Representative micro-photographs show relative expression of NFH (A-fibers) and pgp9.5 (all fibers) positive fibers in MM, TM, and LPM of adult male marmosets. Blue arrows indicate pgp9.5^+^/NFH^-^ fibers. Pictures from MM, TM and LPM, as well as antibodies used, and corresponding colors are indicated. Scales are presented in each microphotograph.

**Figure 4.**
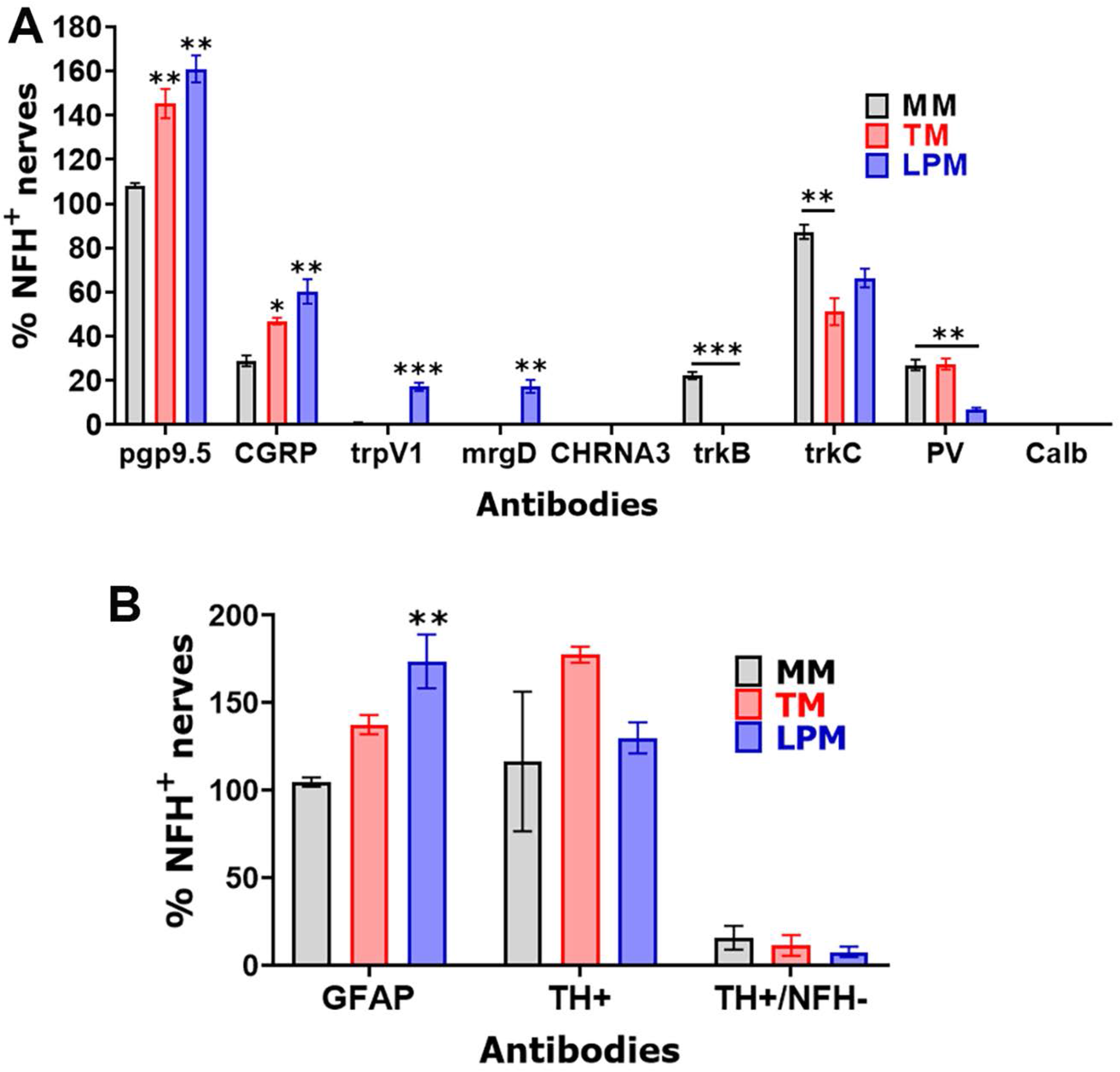
Percentages of different fiber types in MM, TM and LPM of adult male marmosets. (**A**) Baragraphs reflect percentages of marker-positive sensory fibers relative to NFH (A-fibers) in MM, TM, and LPM of adult male marmosets. (**B**) Baragraphs reflect percentages of GFAP^+^ (myelinated) and TH^+^ (sympathetic) fibers relative to NFH (A-fibers) in MM, TM, and LPM of adult male marmosets. X-axis denotes markers and tissue types (i.e. MM, TM and LPM). N=3 for MM and N=2 for TM and LPM.

A-fibers are traditionally considered myelinated, while C-fibers are unmyelinated^31^. Nevertheless, in certain tissues several C-fibers are wrapped by myelin sheath^40^. To examine whether C-fibers are wrapped by myelin sheath in masticatory muscles of adult marmosets, we labeled myelinated fibers with GFAP antibodies. A majority of reports, except few^41^, consider GFAP as a marker for astrocytes, Schwann cells and satellite glial cells. We additionally validated GFAP antibodies in marmosets trigeminal ganglia (TG) (*Suppl Fig 1*)^42^. All NFH^+^ nerves in MM were labeled with GFAP, but about 4% nerves were GFAP^+^/NFH^-^ (*Figs 4B, 5*). NFH^+^ nerves in TM and LPM were also co-labeled with GFAP (*Fig 5*). Proportions of GFAP^+^/NFH^-^ fibers in TM and, especially LPM were significantly higher compared to MM (*Figs 4B, 4*; 1-way ANOVA; F (2, 4) = 20.03; P=0.008; n=2-3). We found that TM and LPM contained ≈28% and ≈43% GFAP^+^/NFH^-^ fibers, respectively (*Fig 4B*). Altogether, the masticatory muscles contain a subset (lowest percentage in MM and highest in LPM) of myelinated C-fibers. These myelinated C-fibers could represent bundles of nerves wrapped with myelin sheathes, which were observed in dura mater^43^.

**Figure 5.**
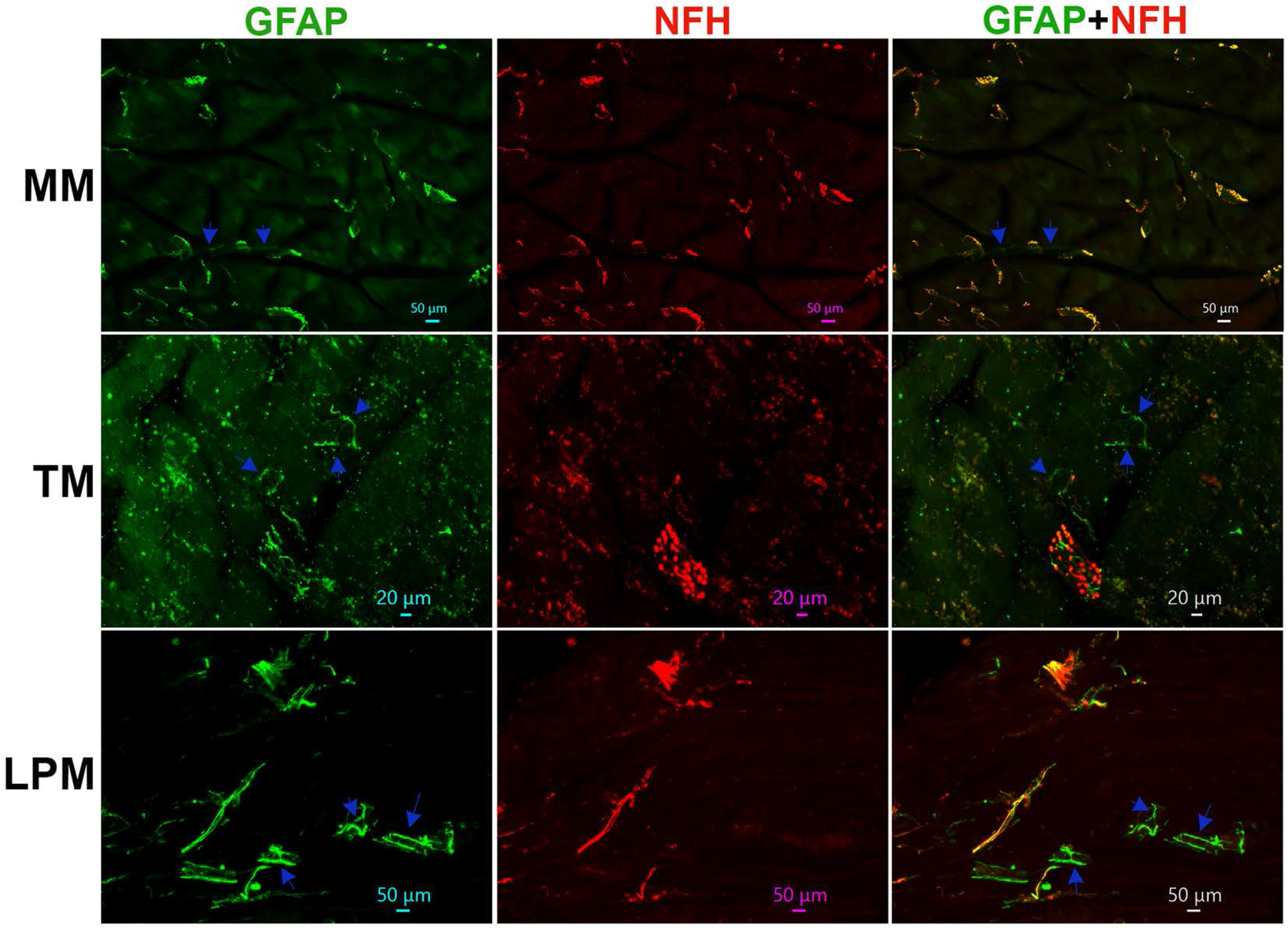
Representations of myelinated fibers in MM, TM and LPM of adult male marmosets. Representative micro-photographs show GFAP myelinated fiber distributions relative to NFH^+^ fibers in MM, TM, and LPM of adult male marmosets. Blue arrows indicate GFAP^+^/NFH^-^ fibers. Pictures from MM, TM and LPM as well as antibodies used and corresponding colors are indicated. Scales are presented in each microphotograph.

### Expression of CGRP^+^ peptidergic nerves in the masticatory muscles

A standard marker for peptidergic nerves and neurons is CGRP^44,45^. We used validated anti-CGRP antibodies, which exhibited strong labeling in a subset of TG neurons in adult male marmosets (*Suppl Fig 1*). Compared to CGRP^+^ signal strength in marmosets TG, CGRP^+^ labeling in masticatory muscle was significantly dimmer (*Fig 6 and Suppl Fig 1*). MM had about 23% of CGRP^+^ nerves, and among them, only ≈3% were CGRP^+^/NFH^-^ (*Figs 4A, 6*). The proportion of CGRP^+^ fibers were higher in TM and particularly in LPM compared to MM (*Figs 4A, 6*; 1-way ANOVA; F (2, 4) = 24.30; P=0.006; n=2-3). Nevertheless, less than half of CGRP^+^ positive fibers were GFAP^+^/NFH^-^ in TM and LPM (*Fig 4A*). These data indicate that peptidergic nerves in the masticatory muscles, especially MM, are predominantly composed of A-fibers.

**Figure 6.**
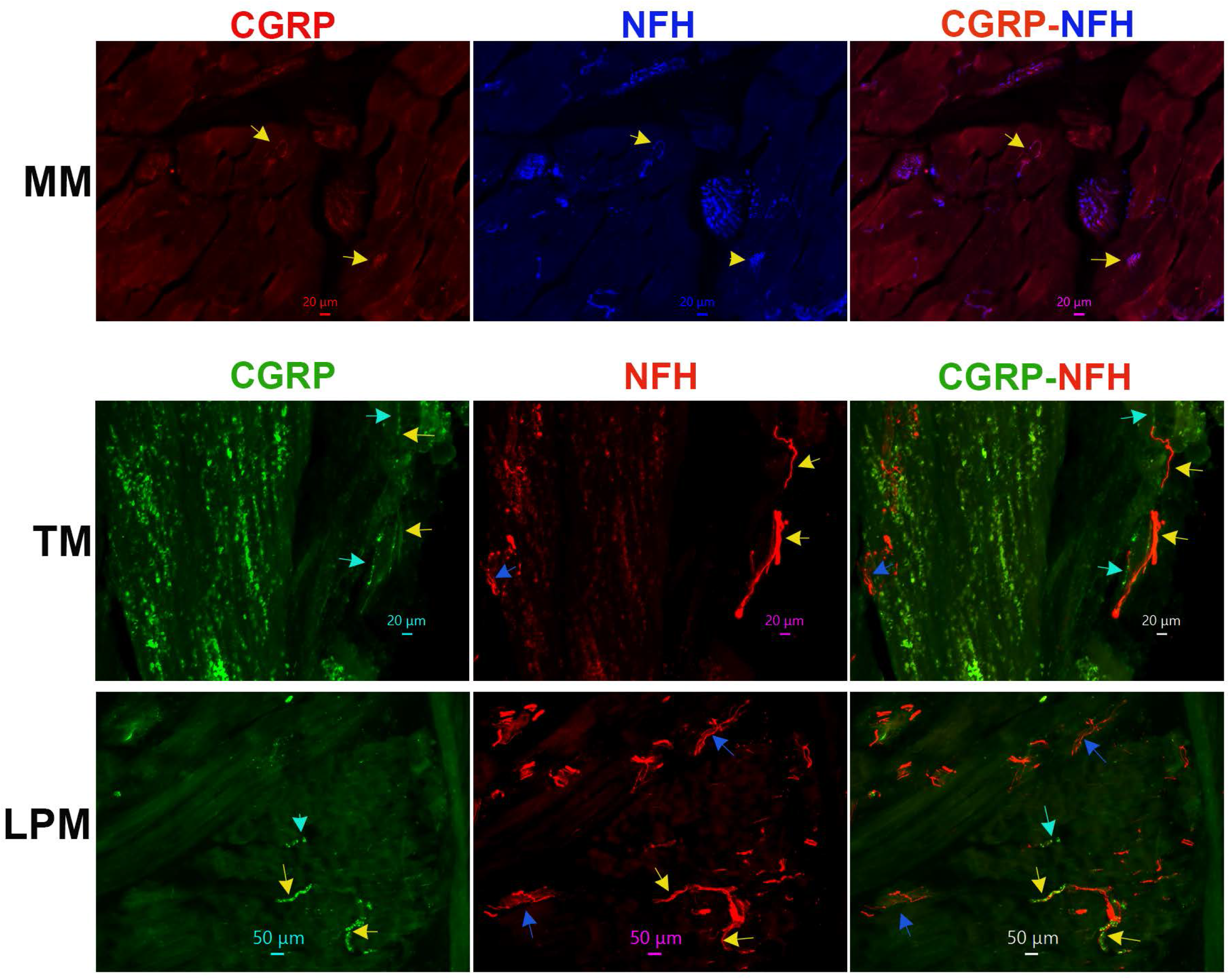
Distribution of peptidergic fibers in MM, TM and LPM of adult male marmosets. Representative micro-photographs show CGRP-positive peptidergic fiber distributions relative to NFH^+^ fibers in MM, TM, and LPM of adult male marmosets. Yellow arrows indicate CGRP^+^/NFH^+^ fibers, cyan arrows show CGRP^+^/NFH^-^ fibers, and blue arrows mark CGRP^-^/NFH^+^ fibers. Pictures from MM, TM and LPM as well as antibodies used and corresponding colors are indicated. Scales are presented in each microphotograph.

### Expression of TrpV1 and MrgprD in the masticatory muscles

Data from *Figures 1-6* imply that masticatory muscles have relatively smaller subset of C-fibers compared to skin and dura mater and are predominantly innervated by A-fibers^43,46,47^. Here, we have evaluated expressions of trpV1, a marker for a subset of C-fiber sensory neurons^44^, and MrgprD, a marker for non-peptidergic C-fiber sensory neurons^44,45^, in the masticatory muscles. TrpV1 and MrgprD antibodies were validated on TG sections, and produced strong labeling in subsets of marmoset TG neurons (*Suppl Fig 1*). As expected from pgp9.5, NFH and CGRP co-labeling, MrgprD fibers were not detected in MM, TM and LPM (*Figs 4A, 8*). Surprisingly, despite capsaicin produced behavioral responses in animals and humans, trpV1^+^ fibers were not identified in MM, TM and LPM (see “Discussion”; *Figs 4A, 7, 8*). Nevertheless, some NFH^+^ nerve bundles within LPM contained both trpV1 and MrgprD labeling (*Figs 4A, 8*). This unusual labeling for trpV1 and MrgprD in LPM could belong to stretches of MN or ATN nerve trunks, which pass through LPM towards the TMJ ligament (see “Discussion”; *Fig 2* schematic). Overall, we found that MM, TM and LPM did not show immunoreactivity for trpV1^+^ or a marker for non-peptidergic sensory fibers.

**Figure 7.**
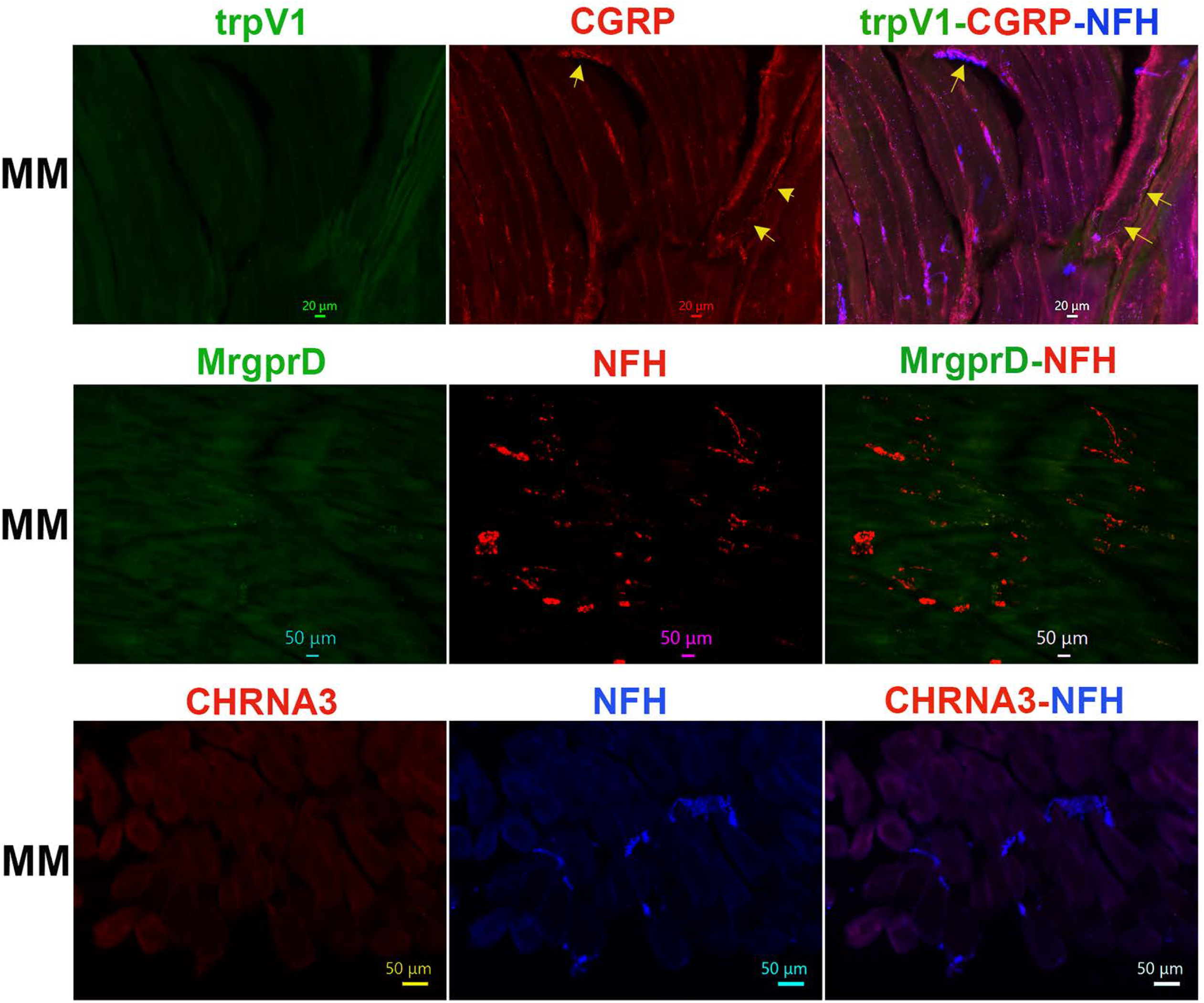
Distribution of trpV1-, MrgprD and CHRNA3-positive fibers in MM of adult male marmosets. Representative micro-photographs show trpV1^+^, mrgprD^+^ non-peptidergic fiber, and CHRNA3^+^, marker for “silent” nociceptors, fiber distributions relative to CGRP^+^ and/or NFH^+^ nerves in MM of adult male marmosets. Yellow arrows in top row panels mark CGRP^+^/NFH^+^/trpV1^-^ fibers. Pictures from MM as well as antibodies used and corresponding colors are indicated. Scales are presented in each microphotograph.

**Figure 8.**
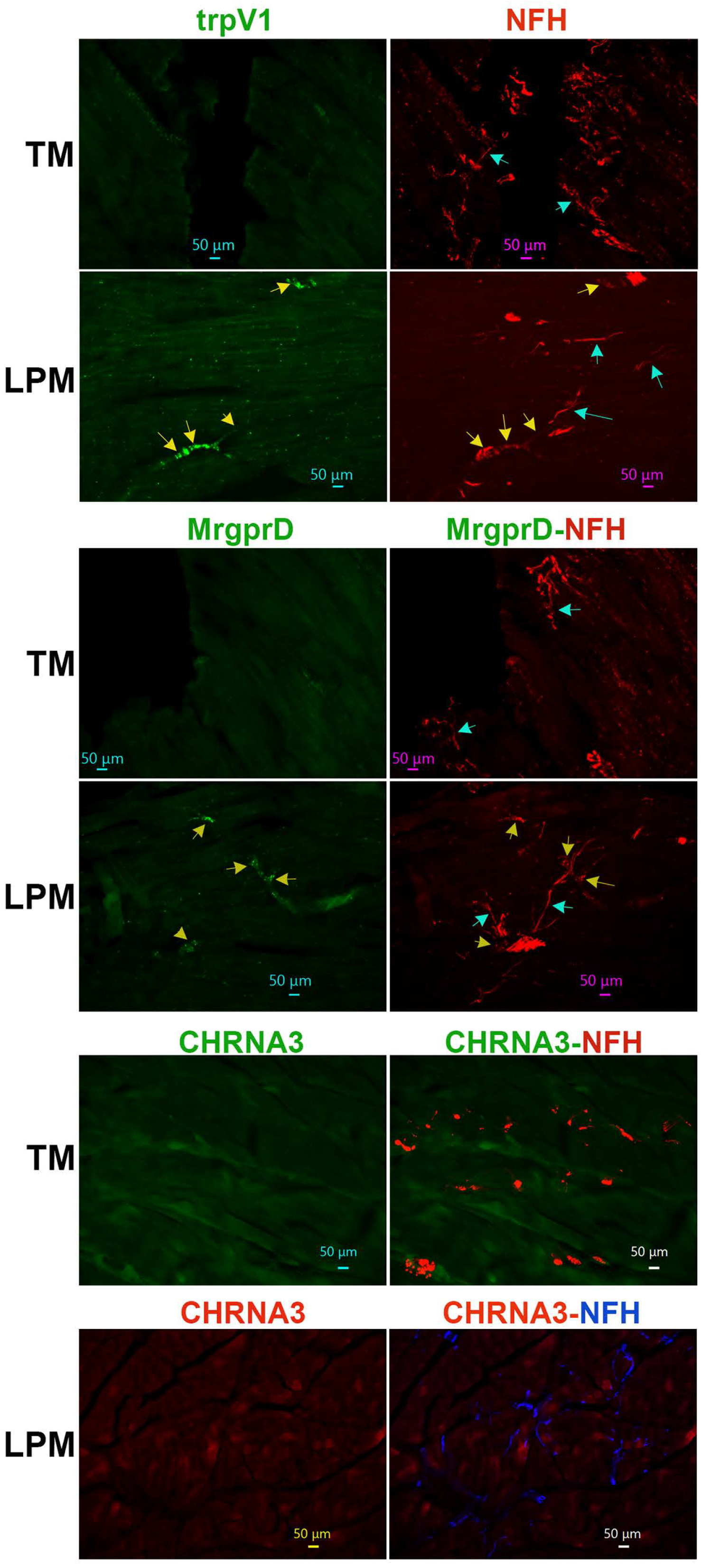
Distribution of trpV1-, MrgprD and CHRNA3-positive fibers in TM and LPM of adult male marmosets. Representative micro-photographs show trpV1^+^, mrgprD^+^ and CHRNA3^+^ fiber distributions relative to NFH^+^ nerves in TM and LPM of adult male marmosets. Yellow arrows (panels for trpV1 and mrgprD) indicate trpV1^+^/NFH^+^ or mrgpr^+^/NFH^+^ fibers, while cyan arrows (panels for trpV1 and mrgprD) show trpV1^-^/NFH^+^ or mrgpr^-^/NFH^+^. Pictures from TM and LPM as well as antibodies used and corresponding colors are indicated. Scales are presented in each microphotograph.

### Expression of a marker for “silent” nociceptors in the masticatory muscles

Certain nociceptors were labeled “silent” nociceptors, since they are insensitive to piezo device stimulation or respond only to very high-threshold *in vitro* or *in vivo* mechanical stimuli. These nociceptors are a subset of C-fibers in DRG and identified by expression of the nicotinic acetylcholine receptor subunit alpha-3 (CHRNA3)^48^. We used validated anti-CHRNA3 antibodies, which show labeling of fibers in the togue of common marmosets (*Suppl Fig 1*), to identify CHRNA3^+^ fibers in masticatory muscles. Unlike NPH tongue, MM, TM and LPM did not express CHRNA3 nerves at a detectable level (*Figs 7, 8*). We also attempted to label marmoset masticatory muscles with several different human PIEZO2 antibodies. They have either did not show any signal or labeling could have been attributed to autofluorescence.

### Expression of A-fiber markers in the masticatory muscles

Single-cell RNA sequencing data and electrophysiology studies on reporter mice indicate that tyrosine hydroxylase (TH), trkB, trkC, parvalbumin (PV) and calbindin-28d (Calb) are markers for DRG low-threshold mechanoreceptor (LTMR) sensory neurons and cutaneous A-fibers^44,45,49,50^. Validated antibodies for trkB, trkC, PV and Calb labeled a subset of TG neurons in marmosets (*Suppl Fig 2*), whereas we observed no TH labeling of neurons or non-neuronal cells in marmosets TG (*Suppl Fig 1*). TrkB as a marker for Aδ-LTMR in skin labeled about 22% of NFH^+^ fibers in MM (*Figs 4A, 9*)^51^. In contrast, TM and LPM did not have detectable trkB labeling (*Figs 4A, 10*; 1-way ANOVA; F (2, 4) = 105.0; P=0.0003; n=2-3). A marker for Aβ-LTMR, trkC, was present on a majority of NFH^+^ fibers in MM (*Figs 4A, 9*)^44,50^. However, trkC was identified only in a subset of A-fibers in TM and LPM (*Fig 10* and *Suppl Fig 3*). The percentages of trkC^+^ A-fibers in TM and LPM were significantly less than those found in the MM (*Figs 4A, 9, 10*; 1-way ANOVA; F (2, 4) = 54.95; P=0.0012; n=2-3). Certain trkC^-^/NFH^+^ and trkC^+^/NFH^+^ fibers were located at the junction of muscle and tendon in TM (*Suppl Fig 3*). PV is another traditional marker for certain Aβ-LTMR and proprioceptors in DRG^44,50^. PV^+^ fibers were present among NFH^+^ nerves in MM and TM, and to a lesser extent in LPM (*Figs 4A, 9, 10*; 1-way ANOVA; F (2, 4) = 24.02; P=0.006; n=2-3). Calb expression was detected in the Aβ-field (aka NF2) sensory neuron type ^44,52,53^. Calb^+^ nerves were not found in MM, TM or LPM (*Figs 4A, 9, 10*). Overall, our data suggest that the masticatory muscles are innervated by A-fibers, including A-LTMR, which are defined by expression patterns of trkB, trkC, Calb and PV. Types of A-fibers are unique for MM, TM and LPM. MM had the highest proportion of A-LTMR among NFH^+^ nerves (i.e. A-fibers).

**Figure 9.**
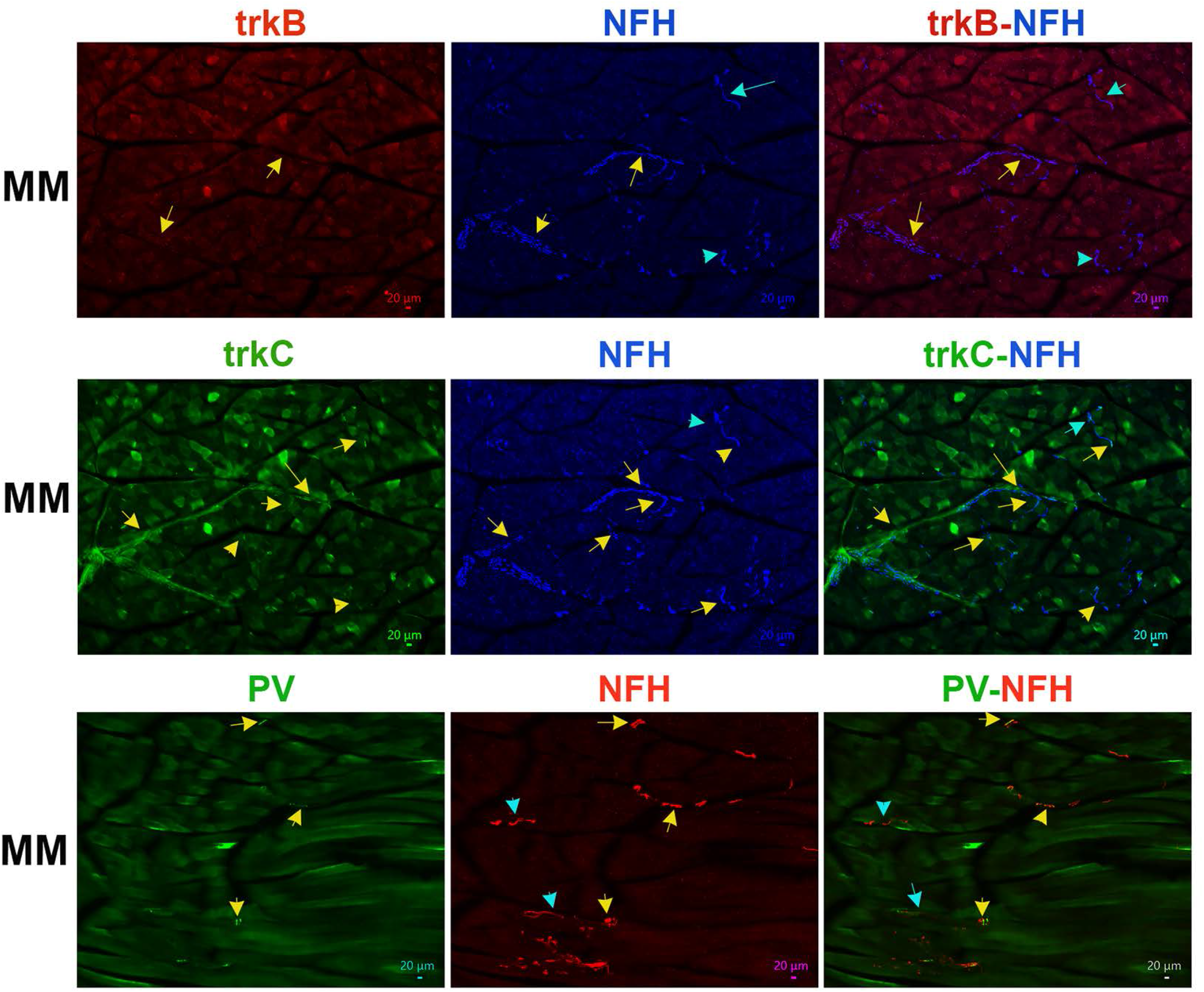
Distribution of trkB, trkC and parvalbumin (PV)-positive fibers in MM of adult male marmosets. Representative micro-photographs show expression of markers for non-nociceptive neurons, trkB, trkC and parvalbumin (PV), in sensory fibers in MM of adult male marmosets. NFH was used to outline all A-fibers in the tissues. Yellow arrows in the top row panels indicate trkB^+^/NFH^+^ fibers, while cyan arrows show trkB^-^/NFH^+^ fibers. Yellow arrows in the middle row panels indicate trkC^+^/NFH^+^ fibers,. Yellow arrows in the bottom row panels indicate PV^+^/NFH^+^ fibers, while cyan arrows show PV^-^/NFH^+^ fibers. Pictures from MM as well as antibodies used and corresponding colors are indicated. Scales are presented in each microphotograph.

**Figure 10.**
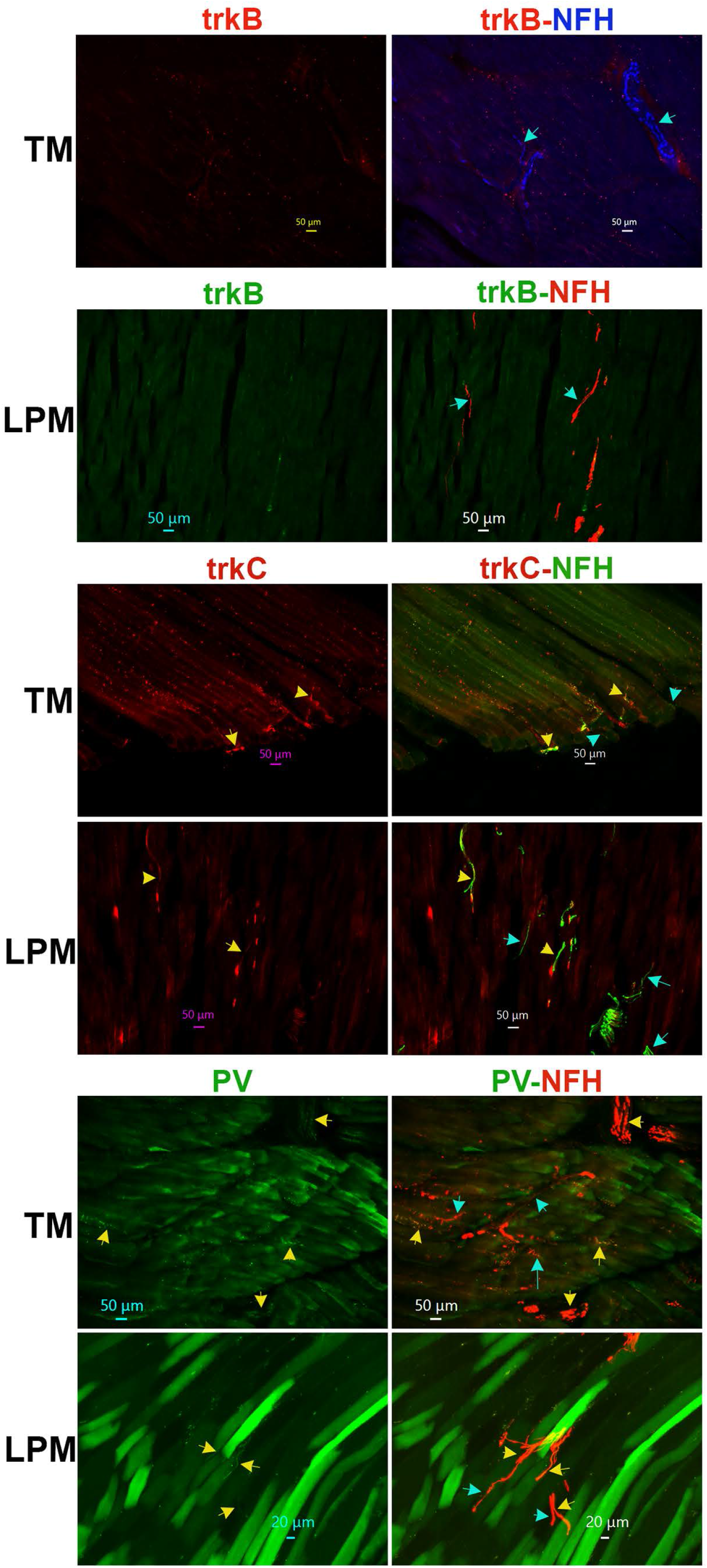
Distribution of trkB, trkC and parvalbumin (PV)-positive fibers in TM and LPM of adult male marmosets. Representative micro-photographs show expression of trkB, trkC and PV sensory fibers in TM and LPM of adult male marmosets. NFH was used to outline all A-fibers in the tissues. Yellow arrows show and trkC^+/^NFH^+^ (middle panels) and PV^+^/NFH^+^ (bottom panels) sensory fibers, while cyan arrows mark trkB^-^/NFH^+^ (top panels), trkC^-/^NFH^+^ (middle panels) and PV^-^/NFH^+^ (bottom panels) sensory fibers. Pictures from TM and LPM as well as antibodies used and corresponding colors are indicated. Scales are presented in each microphotograph.

### Innervation of blood vessels in the masticatory muscles

Muscle ischemia could play an important role in initiation of TMDM. Hence, we investigated innervation of muscle blood vessels. We have used TH as a marker for sympathetic nerves. TH is also a marker for cutaneous C-LTMR^50,54^. However, we could not identify TH^+^ sensory neurons in marmosets TG (*Suppl Fig 1*). Expansive presence of TH^+^ nerves was revealed in the masticatory muscles (*Figs 4B, 11*). The numbers of TH^+^ fibers were comparable to NFH^+^ ones in all studied masticatory muscles (*Figs 4A, 11*; 1-way ANOVA; F (2, 4) = 0.95; P=0.46; n=2-3). A majority of TH^+^ fibers were not co-localized with NFH, but positioned around alpha-smooth muscle actin (α-SMA^+^) blood vessels in MM, TM and LPM (*Figs 4A, 11*; 1-way ANOVA; F (2, 4) = 0.45; P=0.67). Overall, TH^+^ nerves representing sympathetic fibers are located around blood vessels in MM, TM and LPM. Certain portion of sensory NFH^+^ fibers are also in a vicinity of masticatory muscle blood vessels as well as nearby of TH^+^ fibers (*Fig 11*).

**Figure 11.**
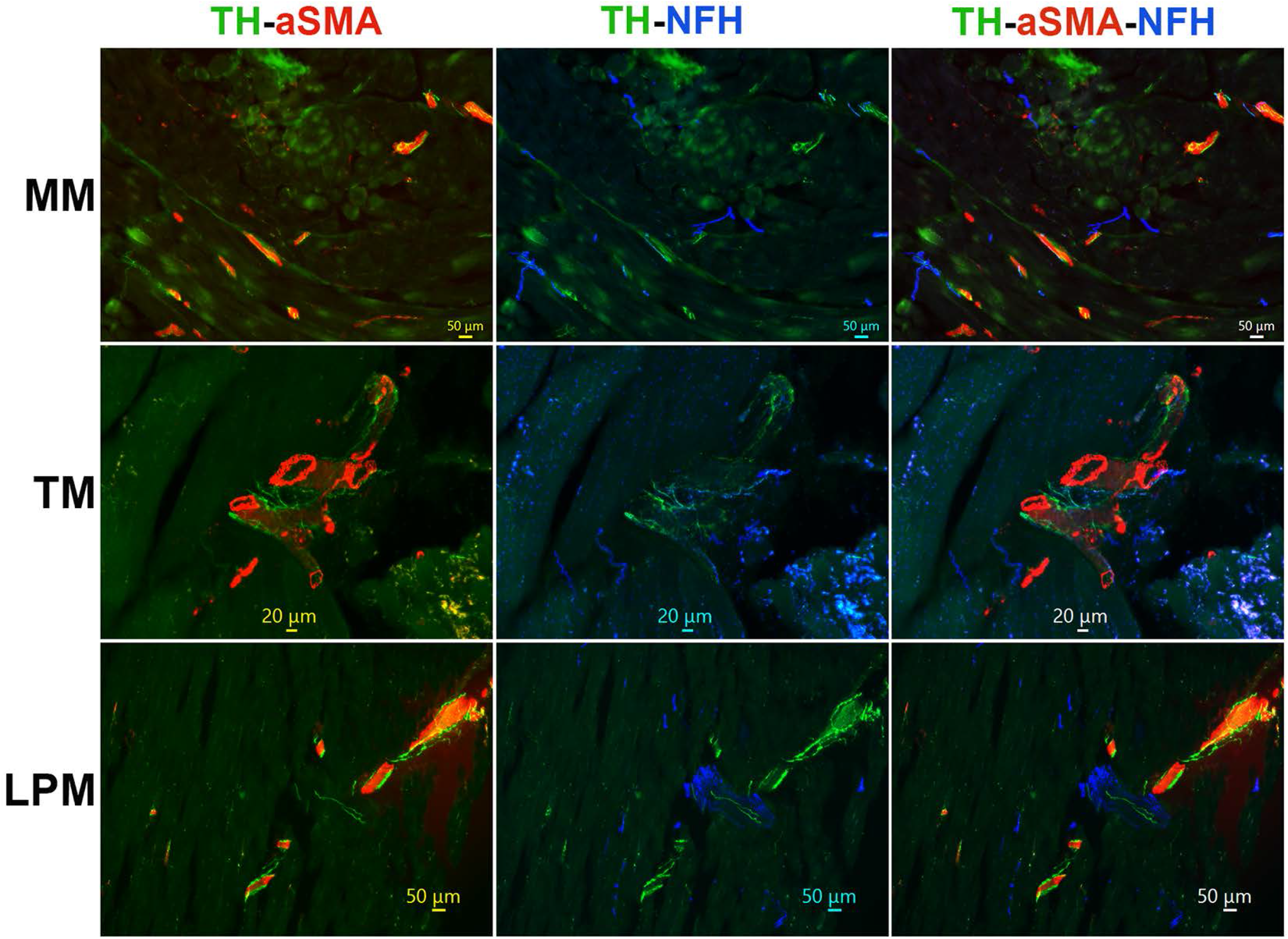
Location of sympathetic fibers in MM, TM and LPM of adult male marmosets. Representative micro-photographs show the location of tyrosine hydroxylase (TH)-positive sympathetic fibers relative to blood vessels outlined with smooth muscle marker alpha-smooth muscle actin (α-SMA) and sensory A-fibers labeled with NFH in MM, TM, and LPM of adult male marmosets. Pictures from MM, TM and LPM as well as antibodies used and corresponding colors are indicated. Scales are presented in each microphotograph.

## Discussion

The etiology and pathogenesis of TMDM is poorly understood^6,55,56^. However, there is an agreement that TMD-M leads to sensitization of nociceptive and non-nociceptive TG sensory neurons innervating the masticatory muscles, and thereby enhancing signal input into the central nervous system and inducing pain^10^. Thus, in order to improve our knowledge about chronic TMDM and possible therapeutic targets, it is essential to characterize sensory nerves in the masticatory muscles. Such information will have higher translatability to clinical application when it is obtained from either humans or animal species that have similar neuroanatomy to humans. Current single-cell RNA sequencing data imply that DRG and TG sensory neurons have similar neuronal clusters^25,52,57^. Nevertheless, it was demonstrated for several tissues that expression patterns for a variety of genes and electrical activity of DRG and TG sensory neurons depends on innervation targets^20-22^. TMDM is prevalent in females (70%) compared to males (30%). Studies on gene expressions in females and males sensory neurons showed that they are different^58-60^. However, sensory neuron types are the same in males and females^24,44,53^. Taken together, this study was conducted on common marmosets, a nonhuman primate species that more closely represents human physiology, genetics, and anatomy than do rodent models. We selected key masticatory muscles, MM, TM and LPM, for this study, since they innervation could be distinct. Studies were conducted only in males, since we aim to investigate type of neurons, but not their transcription profiles.

Masticatory muscles, especially MM, contained a substantially higher proportion of A relatively to C-fibers (*Fig 12*). This innervation pattern is similar to those observed in mouse MM, which is almost exclusively innervated by A-fibers^20^. C-fibers in the masticatory muscles of adult male marmosets are peptidergic, lack non-peptidergic neuronal marker mrgprD and not labeled with trpV1 antibodies. Absence of mrgprD+ and trpV1^+^ immunoreactivities and lack of trpV1-GFP^+^ and mrgprD-TdTom^+^ nerves were also reported in mouse MM^20^. Masticatory muscles also lack the marker for the “silent” nociceptor, CHRNA3^48^. It is surprising that we could not detect trpV1^+^ nerves in primates as well as mouse masticatory muscles^20^. Behavioral experiments on rodents and clinical data demonstrated that intramuscular capsaicin (>10μg) elicits nociception in rodents and pain and crump-like sensation for humans^47,61-64^. One of possible explanations for this difference is that low levels of trpV1 on nerves is sufficient for responses to capsaicin stimulation, while substantially higher expression of trpV1 on fibers is required to reveal immunoreactivity. Another possibility is that trpV1 is expressed on non-neuronal cells surrounding sensory fibers; and capsaicin sensitize nerves indirectly by activating surrounding non-neuronal cells. It is also possible that injected capsaicin defused to a mucosal part, loose areolar and subcutaneous as well as pericranium tissues of facial skin covering muscles, which could contain trpV1^+^ nerves.

**Figure 12.**
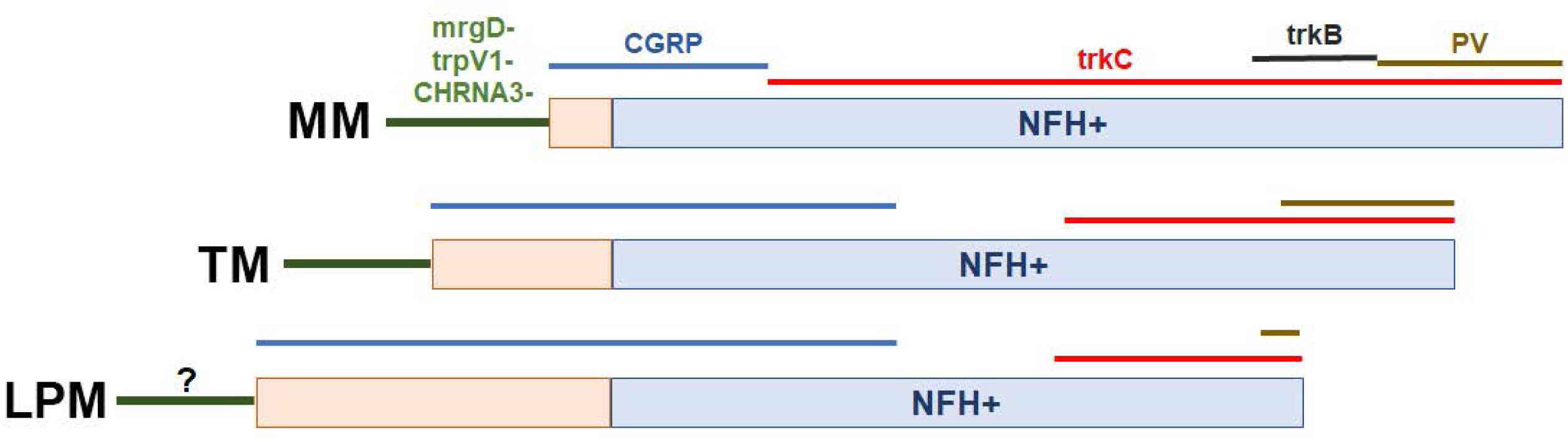
Schematic for sensory neuron subtypes and marker expression in masticatory muscles. Masticatory muscles type (i.e. MM, TM and LPM) are indicated from left. Sensory nerve groups innervating MM, TM or LPM are marked by lines and corresponding marker. *?* indicates unclear presence of mrgprD and trpV1 markers.

We observed possible mrgprD and trpV1 immunoreactivity in stretches of NFH containing nerve trunks in LPM, while mrgprD^+^/NFH^-^ and trpV1^+^/NFH^-^ nerves were absent in LPM. This phenomenon could be explained by the fact that LPM is innervated by a mandibular nerve branch – MM and ATN, which travels through LPM on the way to innervating TMJ ligaments^12^. Therefore trpV1, mrgprD and NFH could belong to such a nerve trunk targeting TMJ. Nevertheless, we do not exclude that this could be non-specific labeling.

C-fibers are traditionally considered un-myelinated fibers. However, it was reported that several C-fibers could compose a nerve trunk, which is wrapped in a myelin sheath^43^. Our data imply that C-fibers in the masticatory muscles could also be wrapped in myelin sheath, which can be detected by labelling with GFAP. Precise organization of C-fibers in these muscles will need to be uncovered using electron microscopy. C-fibers could lose GFAP^+^ sheath during last 30-100μm stretch before innervating targets as they do in dura mater^43^.

Primate TG neurons also lack TH labeling in TG. It may indicate that the masticatory muscles do not have C-LTMR. On other hand, extensive network of TH^+^ fibers, which do not co-localize with NFH fibers was detected around blood vessels in MM, TM and LPM. One possible identity for such TH^+^ fibers are sympathetic nerves.

Comparing our results with data from single-cell RNA sequencing of primate DRG neurons imply that masticatory muscle C-fibers are not similar to PEP1 group containing trpV1. PEP2 and PEP3 sensory neuronal groups are peptidergic with no trpV1 expression^53^. However, PEP2 and PEP3 represent A-fibers (A-HTMR groups) and are similar to mouse CGRP-eta^52,53^. Considering that primate masticatory muscles have NFH^+^ peptidergic fibers, which probably belong to the A-HTMR group ^45,52^ and similar to reported PEP2 and PEP3 clusters in primates (*Fig 12*). Identity of C-fibers in these muscles are not clear.

The remaining A-fibers could be classified as A-LTMR^50^ (*Fig 12*). MM, TM and LPM have no expression of calbindin, which is a marker for NF2 cluster in DRG sensory neurons^44^. The cutaneous Aδ-LTMR marker, TrkB, was found only in MM at low levels (*Fig 12*), and most likely has a distinct function when compared to Aδ-LTMR fibers innervating hair^49,51^. Parvalbumin is a marker for Aβ-LTMR and was present in few nerves of masticatory muscles. Dominant marker for A-LTMR in MM, TM and LPM was shown to be trkC (*Fig 12*). Altogether, innervation of the masticatory muscles in marmosets is clearly distinct from cutaneous fibers and primate DRG neurons. Thus, it is unlikely that A-LTMR in masticatory muscles terminate in specialized cell structures, such as Merkel cells, Messner or Pacinian corpuscles as they do in skin. Moreover, we also noted substantial differences in nerve types between MM, TM and LPM. These differences can be precisely identified only by detailed single-cell RNA sequencing. Nevertheless, there is a similarity between MM innervation in primates versus mice^20^.

The main conclusions of this study are that (1) the nerves innervating marmoset masticatory muscle are unique compared to cutaneous nerves; (2) innervation nerve subgrouping depend on masticatory muscle types; (3) information available thus far suggests that NHP primate and mouse nerves innervating MM could be quite similar; and (4) highest genetic association of PEP1 and NP2 to human pain states^53^ may not apply for pain conditions in the head and neck area, including TMDM.

## Material and methods

References after 60 have been moved to Supplementary information section

### Animals and ethical approval

The reporting in the manuscript follows the recommendations in the ARRIVE guidelines (PLoS Bio 8(6), e1000412,2010). All animal experiments conformed to IASP and APS’s Guiding Principles in the Care and Use of Vertebrate Animals in Research and Training. We also followed guidelines issued by the National Institutes of Health (NIH) and the Society for Neuroscience (SfN) to minimize the number of animals used and their suffering. All animal experiments conformed to protocols approved by the University Texas Health Science Center at San Antonio (UTHSCSA) and Texas Biomedical Research Institute (TBRI, San Antonio, TX) Institutional Animal Care and Use Committee (IACUC). IACUC protocol title is “Plasticity of Lymphotoxin-beta signaling and Orofacial pain in non-human primates”, and numbers are 20200021AR from UTHSCSA and 1821 CJ 0 from TBRI.

For our studies, we collected tissue from 4-11 adult male marmosets for all experiments. Animals were housed at the UTHSCSA or TBRI. Samples for this study were collected opportunistically, including “Tissue Share program in UTHSCSA and TBRI, from animals that were euthanized at IACUC or TBRI approved endpoints on their respective studies. Marmosets used in this study did not have either injury affecting the head and neck area, or systemic infections.

### Tissue collection and Processing

Animals were euthanized by the veterinarians in UTHSCSA and TBRI at defined end points that were determined for each animal. At the point of determined death, superficial and deep heads of MM, TM, superior and inferior heads of LPM with attached TMJ ligament, trigeminal ganglia (TG), hindpaw skin and tongue were dissected and placed in 4% paraformaldehyde (PFA). Tissues were fixed in 4% PFA for 3-4 hours, washed in 3 × 15 min in 0.1M Phosphate Buffer (PB), equilibrated in 10% sucrose in PB at 4°C overnight and cryo-protected and stored in 30% sucrose in PB at −20°C. Tissues were embedded in Neg 50 (Richard Allan Scientific, Kalamazoo, MI); and were cryo-sectioned with the following thickness: TG 20 μm and MM, TM and LPM 30 μm.

### Immunohistochemistry (IHC)

Immunostaining was performed as described previously^65,66^. Briefly, sections were blocked in 4% normal donkey serum (Sigma, St. Louis, MO), 2% bovine gamma-globulin (Sigma-Aldrich, St. Louis, MO) and 0.3% Triton X-100 (Fisher Scientific) in 0.1M PBS for 90 min at RT. Next, tissue sections were incubated for 24-36 hours at RT with primary antibodies. Sections were washed with PBS from unbound primary antibodies, blocked, and incubated for 90 min at RT with appropriate fluorophore conjugated secondary antibodies (Jackson Immuno-Research, West Grove, PA, USA). Finally, tissue sections were washed for 3 x 5 minutes with 0.1M PBS and 2 x 5 minutes in diH_2_O, air-dried, and covered with Vectashield Antifade Mounting Medium (Vectorlabs, Burlingame, CA, USA). In this study, the following previously characterized primary antibodies were used: anti-PGP9.5 rabbit polyclonal (Millipore-Sigma; Burlington, NJ, catalog AB1761-I; 1:300)^67^; anti-Neurofilament H (NF-H) chicken polyclonal (BioLegend; San Diego, CA; catalog 822601; 1:1000)^68^; anti-CGRP guinea pig polyclonal (Synaptic Systems; Goettingen, Germany; catalog 414 004; 1:200)^69^; anti-TRPV1 rabbit polyclonal (Novus Biologicals; Centennial, CO; catalog NBP1-71774SS; 1:200)^70^; anti-mrgprD rabbit polyclonal (Alamone Lab; catalog ASR-031; 1:200)^20,71^; anti-CHRNA3 rabbit polyclonal (Bioss; catalog BS-6455R; 1:200)^72^; anti-tyrosine hydroxylase (TH) chicken polyclonal (Neuromics; Bloomington, MN; catalog CH23006; 1:300)^73^; anti-smooth muscle actin (α-SMA) Cy3-conjugated rabbit polyclonal antibody (Millipore-Sigma; catalog C6198; 1:500); anti-trkC rabbit polyclonal (Aviva Systems Biology, San Diego, CA; catalog ARP51318_P050; 1:200); anti-trkB goat polyclonal (R&D systems; AF1494; 1:200)^74^; anti-parvalbumin rabbit polyclonal (Novus Biologicals; catalogue NB120-11427SS; 1:200)^75^; anti-calbindin D28k rabbit polyclonal (Synaptic Systems; catalogue 214 011; 1:200)^76^.

### Counting of Fibers and Neurons

Images were acquired using a Keyence BZ-X810 all-in-one microscope (Itasca, IL, USA) in the “sectioning” mode. Images were acquired with a 2×, 10× or 20× objective. Settings were determined in such way that no-primary control did not show any positive signal. Then, images were taken using these fixed acquisition parameters across all groups. Control IHC was performed on tissue sections processed as described but either lacking primary antibodies or lacking primary and secondary antibodies. Z-stack IHC images were obtained from 3-5 independent tissue sections from 2-3 primates/isolations. Mean values from these 3-5 sections generated from an isolation represented data for a biological replicate. Thus, these 2-3 primates/ isolations represent the biological replicates (i.e. *n*). Neuronal fibers were counted manually using Image J software to obtain approximate estimation of the numbers of peripheral nerve types innervating muscle tissues. Bundles of fibers grouped together that couldn’t be visually separated were also counted as 1. The nerve fiber types were identified by expressions of specific markers.

### Statistical Analyses

GraphPad Prism 8.0 (GraphPad, La Jolla, CA) was used for statistical analysis. Data are presented as mean ± standard error of mean (SEM). Differences between groups were assessed by chi-square analysis with Fisher’s exact test, unpaired t-test, or 1-way ANOVA with Bonferroni’s post-hoc test. A difference is accepted as statistically significant when p<0.05. Interaction F ratios, and the associated p values are reported.

## Acknowledgements

We would like to thank Dr. Christopher Chen (Texas Biomedical Research Institute, San Antonio, TX) for his advice in working with primate tissues, Dr. Corina Ross in helping obtain primate tissues via the tissue share program at Southwest National Primate Research Center (SNPRC), San Antonio, TX, the veterinarians of SNPRC with help in dissection of tissues, and Mrs. Qun Li and Mrs. Fang-Mei Chang for validation of trpV1 and CGRP antibodies for human/primate tissues.

## Author contribution

A.H.H and K.A.L. conducted a majority of experiments and analysis; S.B., J.M., J.M. and M.T. performed certain experiments and analysis; S.B., J.M., M.T. contributed to tissues preparation; T.M.C., A.B.S. and D.P. performed tissue isolation; D.P., S.R. and A.N.A. analyzed data, edited draft and the final version of the manuscript; A.N.A. designed and directed the project, wrote the first draft of the manuscript; and prepared the final version of the manuscript.

## Additional Information

The authors declare that they have no known competing financial interests or personal relationships that could influence the work reported in this paper.

