## Supplementary text and figures for "Sensory innervation of masseter, temporal and lateral pterygoid muscles in common marmosets"

### Legends to Supplementary Figures:

**Supplementary Figure 1**. ***Positive control experiments with antibodies for nociceptors, "small" sensory neurons, and glia***.

*Top-row panels* show expression of CGRP (peptidergic neurons) and trpV1 in TG of adult male marmoset. *Middle row panels* show expression of GFAP (pan glial marker), mrgprD (a marker for non-peptidergic neurons) and tyrosine hydroxylase (TH; a marker for C-fiber low threshold mechanoreceptor (C-LTMR)) in in TG of adult male marmoset. *Bottom row panel* exhibits location of CHRNA3 (a marker of "silent" nociceptors in DRG) in tongue of adult male marmosets. Yellow arrows mark CHRNA3$^{+}$ fibers in a marmoset tongue section labeled with CHRNA3 and NFH. Antibodies used and corresponding colors are indicated. Scales are presented in microphotographs from top two rows.

**Supplementary Figure 2**. ***Positive control experiments with antibodies for "large, non-nociceptive neurons.***

*Top-row panels* show expression of trkB (a marker for Aδ-LTMR in DRG) relative to CGRP$^{+}$ neurons in marmoset TG, and Calbindin-1 (Calb; a marker for subset of Aβ-LTMR in DRG) in marmoset TG. *Bottom-row panels* present expressions of trkC (a marker for Aβ-LTMR in DRG) in marmoset TG, and parvalbumin (PV; a marker for a subset of Aβ-LTMR in DRG) relative to CGRP$^{+}$ neurons in marmoset TG. Antibodies used and corresponding colors are indicated. Scales are presented in microphotographs for trkC and Calb.

**Supplementary Figure 3**. ***TrkC fiber expression in TM.***

This is an example of trkC$^{+}$/NFH$^{+}$ (cyan arrows) and trkC$^{-}$/NFH$^{+}$ (yellow arrows) sensory fiber expression in TM muscle and tendon tissues. Antibodies used and corresponding colors are indicated. Scales are presented in microphotographs.

*Top-row panels* show expressions of trkB (a marker for Aδ-LTMR in DRG) relatively to CGRP$^+$ neurons in marmoset TG, and Calbindin-1 (Calb; a marker for subset of Aβ-LTMR in DRG) in marmoset TG.

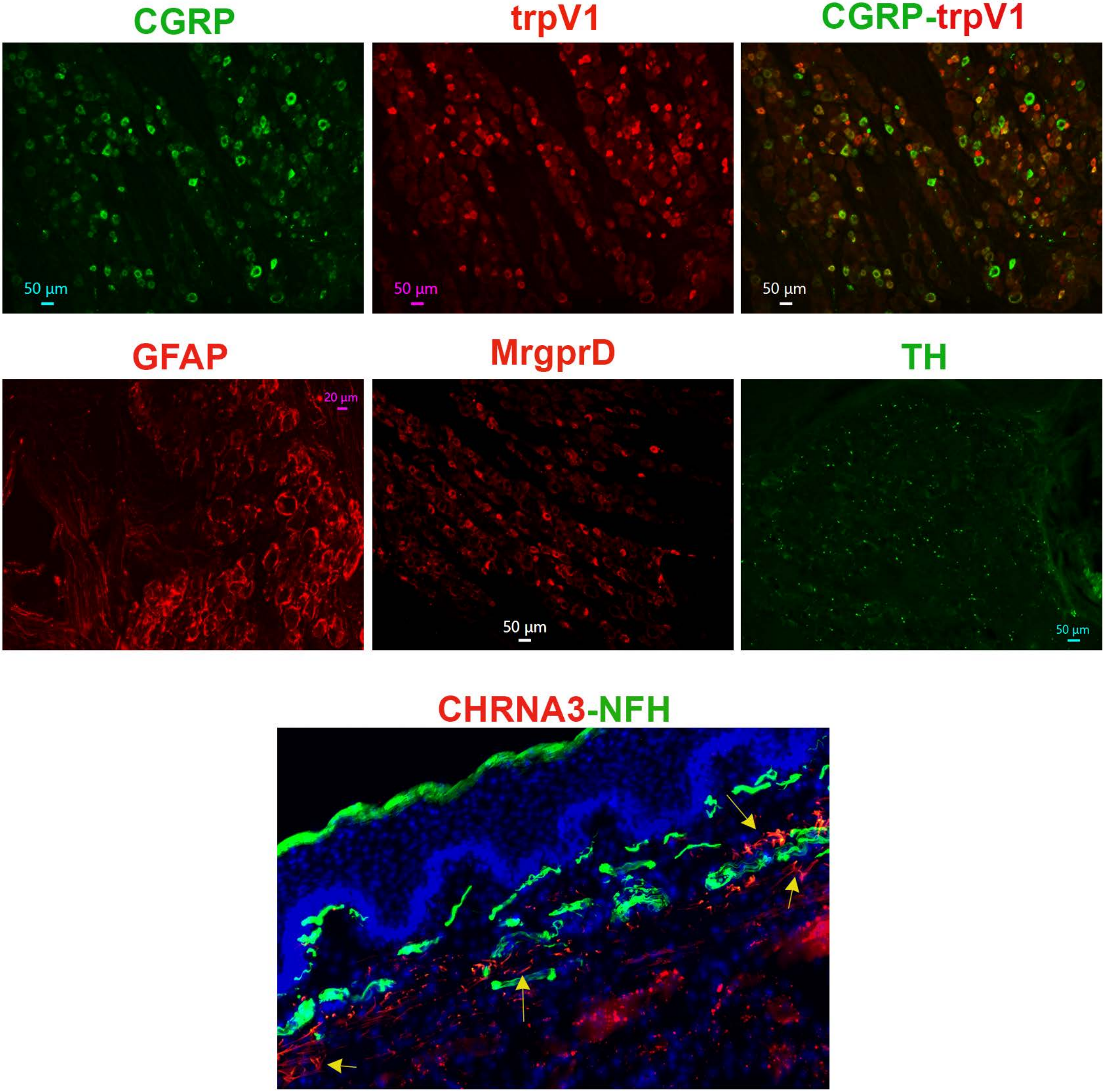

CGRP
trpV1
CGRP-trpV1
50 µm
50 µm
50 µm
GFAP
MrgprD
TH
20 µm
50 µm
50 µm
CHRNA3-NFH

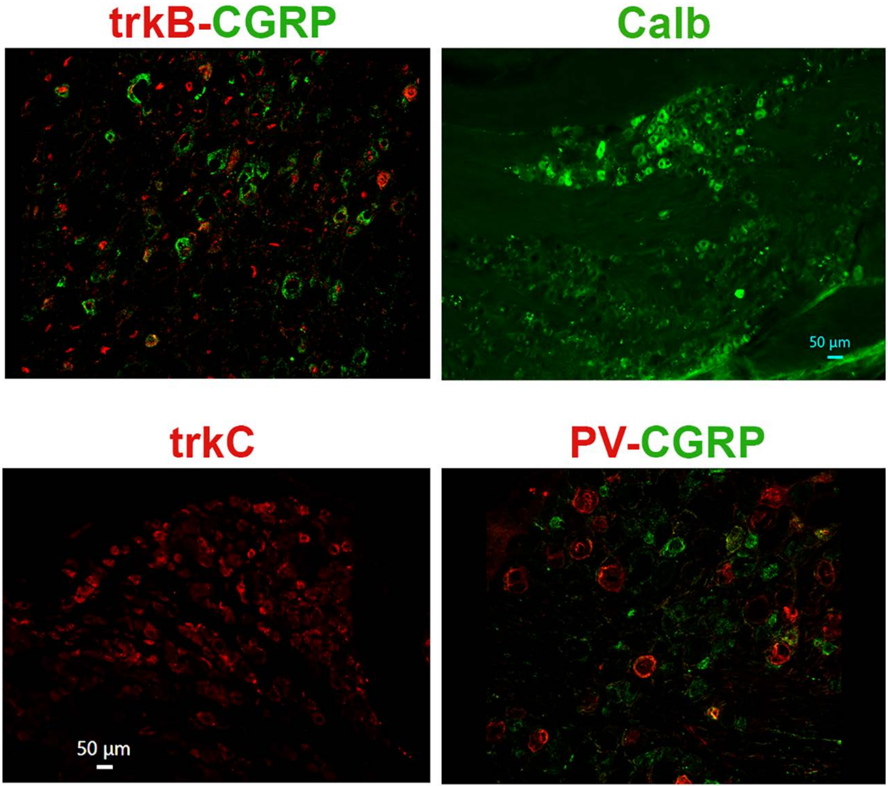
trkB-CGRP
Calb
50 µm
trkC
50 µm
PV-CGRP

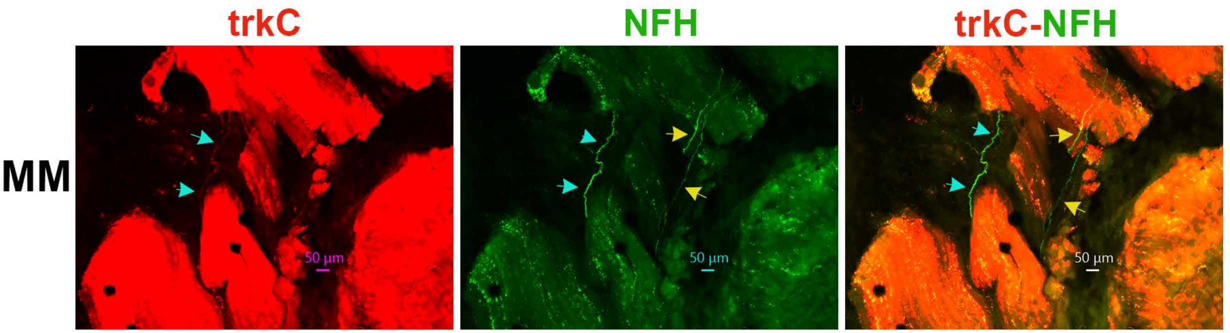
trkC
NFH
trkC-NFH
MM
50 µm
50 µm
50 µm
